# Proteome constrained metabolic modeling of *Sus scrofa* muscle stem cells for cultured meat production

**DOI:** 10.1101/2025.09.30.679571

**Authors:** Sizhe Qiu, Eliska Kratochvilova, Wei E. Huang, Zhanfeng Cui, Tom Agnew, Aidong Yang, Hua Ye

## Abstract

Cultured meat has recently emerged as a sustainable alternative to the traditional livestock farming and gained attention as a promising future protein source. Herein, the *Sus scrofa* muscle stem cell is a commonly used cell source in the cell proliferation step of cultured meat production. However, a major bottleneck of large-scale cultivation is the inhibition by secreted and accumulated lactate and ammonium in the process of *S. scrofa* cell proliferation. To simulate the growth and metabolism of *S. scrofa* muscle stem cells under different lactate and ammonium concentrations, this study constructed the first proteome constrained metabolic model for the core metabolism of *S. scrofa* muscle stem cells, pcPigGEM2025. The relationship of lactate and ammonium levels with cellular metabolism was derived from growth and metabolomics data of two culture conditions with low and high initial ammonium concentrations, and then incorporated into metabolic flux simulation. Metabolic flux simulations for experimental conditions, along with perturbation simulations considering stressed non-growth associated maintenance and oxygen supply, demonstrated that pcPigGEM2025 could effectively characterize the response of the *S. scrofa* muscle stem cell’s growth and metabolism to varying environmental conditions, shedding light on model-aided control and optimization of the cultured meat production process.

**Graphical abstract:** 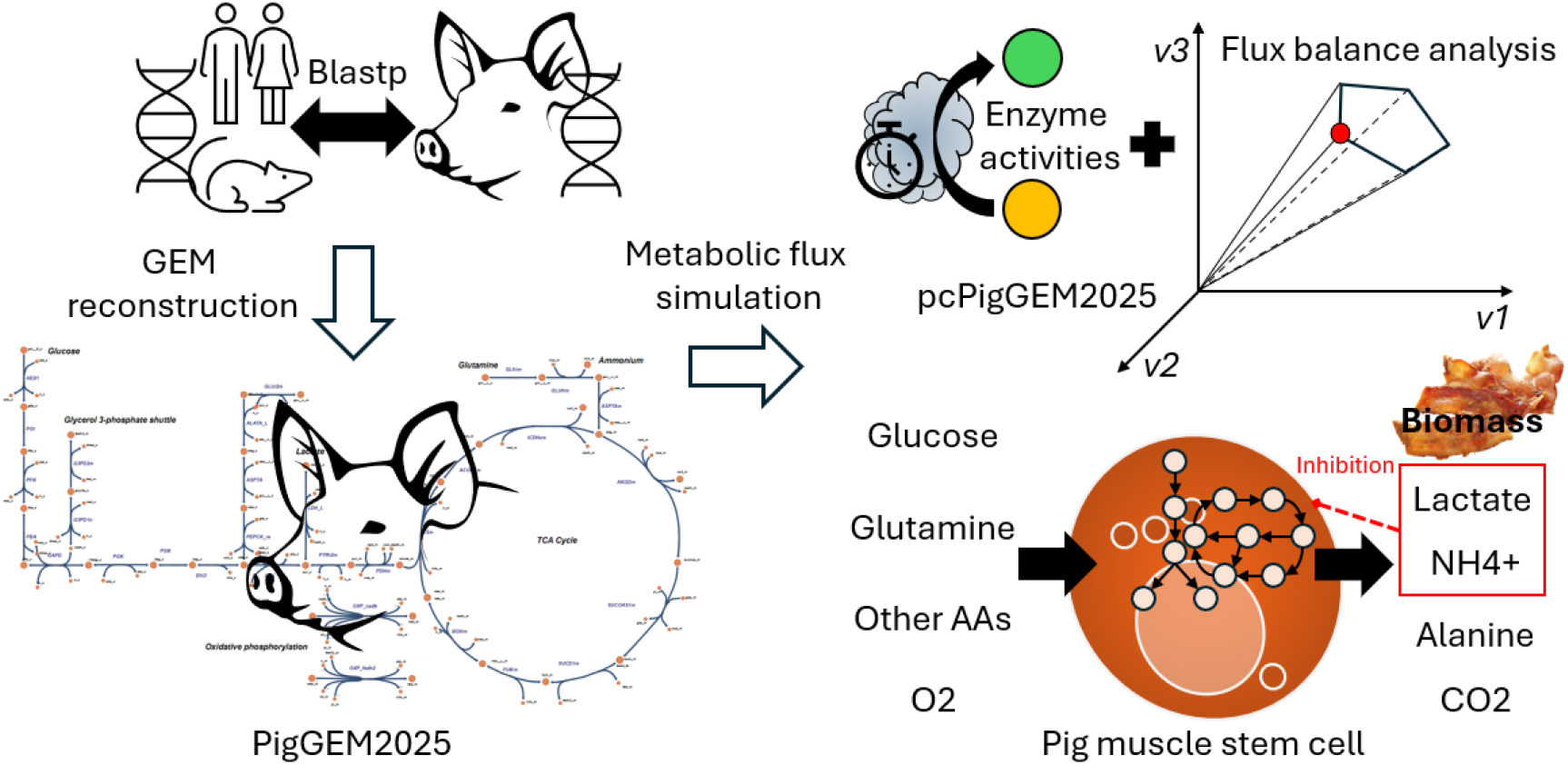

**Highlights:**

1. The first proteome constrained metabolic model was built for *S. scrofa* myoblasts.
2. This model effectively simulated myoblast metabolism under lactate and NH4+ stress.
3. Perturbation simulations showed that this model could also account for other stress.
4. This model enables *in-silico* control and optimization of cultured meat production.

## 1. Introduction

With the rapidly increasing global consumption of meat (Godfray et al., 2018), the traditional livestock farming has placed a growing burden on the environment (Tullo et al., 2019), such as increased land demand, carbon dioxide emission (Caro et al., 2014), and water pollution (Yang et al., 2020). Consequently, cultured meat production from mammalian stem cells has been considered as a sustainable alternative, and the *Sus scrofa* (pig) muscle stem cell is one of the most commonly used cell sources (M. Li et al., 2022; Post, 2012; Zhu et al., 2022). In cultivated pork production, the cell proliferation (biomass production) in the bioreactor is the determining step of cultured meat yield, but the inhibition of cellular growth by secreted and accumulated lactate and ammonium (NH4+) becomes the bottleneck (Chen et al., 2009; Hassell et al., 1991; Ryll et al., 1994; Slivac et al., 2010). As major products of central carbon metabolism and amino acid metabolism, lactate and NH4+ in mammalian cell culture limits cellular growth in a feed-back mode. Therefore, a predictive model of the *S. scrofa* muscle stem cell’s metabolism is desirable to characterize its interactions with the growth environment in the bioreactor for the quantitative analysis and control of cultured meat production process.

Regarding cellular metabolism, flux balance analysis (FBA) with genome scale metabolic models (GEMs) is a widely-used computational method to predict metabolic fluxes under different conditions (Orth et al., 2010). Compared to differential equation-based metabolic models, FBA offers advantages in interpretability and computational efficiency (Qiu et al., 2023a). In recent years, researchers have already built GEMs for several well-studied animal species (Wang et al., 2021). The representative ones are iCHO for Chinese hamster ovary (CHO) cells (Hefzi et al., 2016), RECON1 and RECON-3D for human metabolism (Duarte et al., 2007), and BtaSBML2986 for bovine satellite cells in cultured meat production (Lee et al., 2024). Hence, GEM reconstruction is a feasible approach to analyze the metabolism of *S. scrofa* muscle stem cells. With respect to metabolic flux simulation, Nolan and Lee, 2011 combined differential equation-based kinetics and FBA to dynamically model CHO cell metabolism with changing metabolite concentrations (Nolan and Lee, 2011); Yeo et al., 2020 integrated enzyme capacity constraints into iCHO to achieve accurate flux prediction (Yeo et al., 2020). However, none of these models has incorporated the influence of environmental factors (e.g., the accumulation of lactate) which exist in actual bioprocesses.

With the objective to simulate *S. scrofa* muscle stem cell proliferation under different lactate and NH4+ levels, this study attempted to construct the first GEM of *S. scrofa*’s core metabolism, PigGEM2025, and the first proteome constrained FBA model for *S. scrofa* muscle stem cells, pcPigGEM2025. By approximating the inhibitory effects of lactate and NH4+ using measured growth and extracellular metabolomics data, this study managed to incorporate major environmental stress factors in *S. scrofa* muscle stem cell culture into metabolic flux simulation. Subsequently, the accuracy of pcPigGEM2025 was evaluated through validation against experimental data. Overall, this study intended to present pcPigGEM2025 as a metabolic modeling framework with the potential to be a model-aided process engineering tool for controlling and optimizing cultured meat production.

## 2. Materials and methods

### 2.1 Culture conditions of *S. scrofa* muscle stem cells

Genetically engineered, immortalised, suspension-adapted skeletal-muscle derived stem cells (*S. scrofa*, obtained from Ivy Farm Technologies, Oxford, UK, under a material transfer agreement) were cultured in SR7 medium (formulation by Ivy Farm Technologies, Oxford, UK) supplemented with 1% (v/v) penicillin–streptomycin. The formulation of SR7 medium is based on DMEM/F-12 supplemented with amino acids, lipid concentrate, nucleosides, AlbuMAX, poloxamer 188, anti-clumping agent, fetal bovine serum (FBS) growth factors and others. The exact composition of SR7 medium and the detailed cell culture handling protocols were provided by Ivy Farm Technologies and are proprietary to the company.

Stem cell cultures were maintained at 37 °C in a humidified incubator with 5% CO_2_, in baffled Erlenmeyer flasks with vented filter lids, with the working volume set to one-fifth of flask capacity, and agitated on an orbital shaker (SSM1, Stuart, UK) at 140 rpm for cultures ≤50 mL and 120 rpm for cultures ≥100 mL. For the low initial NH4+ culture experiment, cells at passage 59 were grown in 50 mL SR7 medium at 140 rpm. On day 3, the pre-stationary cultures were seeded into three flasks containing 110 mL medium at a density of 1.0 × 10^5^ cells/mL and incubated at 120 rpm. For the high initial NH4+ culture experiment, SR7 medium was supplemented with ammonium chloride (NH4Cl) to a final concentration of 3.5 mM. Cells at passage 64 (two cultures of 25 mL each, 140 rpm), likewise in pre-stationary phase, were seeded on day 3 into three flasks containing 110 mL medium at a density of 1.0 × 10^5^ cells/mL and incubated at 120 rpm.

At every sampling point, 1 mL samples were withdrawn from each flask and centrifuged at 600 × g for 5 min at room temperature. The resulting cell pellets were treated with TrypLE (Thermo Fisher Scientific, UK) at 37 °C for 5 min, neutralised with fresh medium to the original volume, and mixed 1:1 with 0.4% Trypan Blue Stain (Thermo Fisher Scientific, UK). Cell counts were obtained using an automated cell counter (Countess™, Thermo Fisher Scientific, UK), with each biological replicate value representing the mean of at least two independent technical measurements.

### 2.2 Metabolomics analysis of *S. scrofa* muscle stem cell culture

Following centrifugation, supernatants were collected for metabolomics analysis. In the low initial NH4+ culture experiment, samples were rapidly frozen and stored either overnight at −80 °C before transferred to liquid nitrogen, or snap-frozen directly in liquid nitrogen. In the high initial NH4+ culture experiment, samples were frozen directly at −80 °C. Freezing protocols were selected based on equipment availability. All samples were stored for no longer than one week prior to metabolomics analysis. The analytes quantified in this study (glucose, lactate, glutamine, NH□□) are generally considered chemically stable under frozen storage.

Glucose, lactate, glutamine and NH4+ concentrations were measured using a Nova Flex 2 analyser (Nova Biomedical, UK). The pH was measured at room temperature using a glass combination pH electrode (InLab Micro, Mettler Toledo, UK), calibrated before each measurement session. For both low and high initial NH4+ level conditions, the pH was maintained to be around 7.5, a slightly alkaline pH (**see Figure S1 in SI**). For each metabolite concentration measurement, there were three biological replicates (three independent flasks). The analytical workflow was adapted from the metabolomics profiling of CHO cells (Széliová et al., 2020), with modifications for *S. scrofa* myoblasts. Cell-specific consumption and production rates of glucose, lactate, glutamine and NH4+, and oxygen uptake rates were computed using measured metabolite concentrations and cell counts at different time points (**see section S1**.**1 and Figure S2 in SI**).

### 2.3 Genome-scale metabolic model reconstruction of *S. scrofa* muscle stem cell

To build the genome-scale metabolic model (GEM) of the *S. scrofa* muscle stem cell, namely PigGEM2025 (https://github.com/SizheQiu/PigGEM2025), bidirectional BLASTp (Camacho et al., 2009) was used to find homologous proteins coding for metabolic reactions and then curate gene-protein-reaction rules (GPRs) from template GEMs. RECON1 (Homo sapiens) (Duarte et al., 2007) and iCHO (Cricetulus griseus) (Hefzi et al., 2016) were used as template genome-scale metabolic models (GEMs). The template genomes of *S. scrofa, H. sapiens*, and *C. griseus* used in this study were GCF_000003025.6, GCF_000001405.33, and GCF_003668045.3, respectively. For bidirectional BLASTp, the threshold of e-value was 1e-6. With matched homologous proteins, metabolic reactions were curated based on the boolean computation of GPRs (e.g., GPR=“A or B”, the existence of either gene A or B will include the reaction; GPR=“A and B”, the reaction can only be included if both gene A and B exist). Boundary reactions were added for all metabolites in the extracellular space. The biomass formation reaction (default objective function) was modified from the one in iCHO, based on the experimental measurement of *S. scrofa*’s cellular biomass composition (**see section S1**.**2 in SI**). For metabolic network gap-filling, transportation reactions of organelles and pseudo biosynthetic reactions with no GPR associated were manually added from RECON1 and iCHO to ensure all components in the biomass composition can be produced in FBA. Redundant metabolites and reactions were manually removed.

Next, reactions in the subsystems of glycolysis, pentose phosphate pathway, tricarboxylic acid (TCA) cycle, oxidative phosphorylation, amino acid metabolism, urea cycle, nucleotide metabolism, and lipid synthesis were collected to build a GEM of core cellular metabolism. Metabolites and reactions in the compartments of nucleus, peroxisome/glyoxysome, lysosome, and endoplasmic reticulum were removed. The quality of the finalized core GEM of the *S. scrofa* muscle stem cell was assessed by MEMOTE (Lieven et al., 2020).

### 2.4 Proteome constrained flux balance analysis of *S. scrofa* muscle stem cells

To tighten the metabolic flux solution space, this study integrated proteome constraints of reactions into conventional FBA (Orth et al., 2010) to build a proteome constrained FBA model (Mori et al., 2016) for *S. scrofa* muscle stem cells, namely, pcPigGEM2025. COBRApy (Ebrahim et al., 2013) was used to perform FBA. The objective function was maximization of the growth rate (*μ*), and the default constraint was mass conservation (Eq. 1). *S* is the stoichiometric matrix of the metabolic reaction network, and *v* represents the vector of reaction fluxes (Eq. 1). The reaction flux 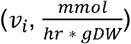 was constrained by the enzyme activity, 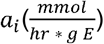 (Eq. 2). Enzyme activity values were obtained from BRENDA (Chang et al., 2021) and SABIO-RK (Wittig et al., 2018). The proteome of *S. scrofa* was divided into five sectors: C sector for carbon catabolism, E sector for biosynthesis of amino acids, lipids, nucleotides, and polysaccharides, A sector for anabolism (biomass formation), T sector for transporter proteins, and NGAM sector for non-growth associated maintenance. *ϕ*_*j*_ represented the mass fraction of the sector *j* for *j = A, C, T, E, NGAM. P*_*TOT*_ was the total mass of the proteome normalized to 1 gDW of biomass 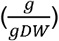 (**see Table S3 in SI**), and the upper bound of the summation of all five sector fractions was 100% (Eq. 3 and 4). Eq. 5-7 were integrated into FBA to model the inhibitory effects of lactate and NH4+.*ub*_*glucose*_, *ub*_*glutamine*,_ and 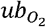 were upper bounds of uptake fluxes of glucose, glutamine, and oxygen. Eq.6 was also applied to the upper bounds of uptake fluxes of other amino acids. Eq.5 was a piecewise function: if 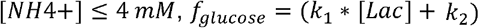; if 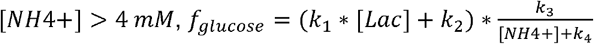. Exponential decay and the Haldane-type inhibition term (Kang et al., 2016) were also tried for Eq. 5, but the performance was inferior to the piecewise function (**see Table S8 in SI**). Eq.6 and Eq.7 were both in the form of 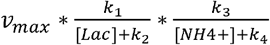. The parameters in Eq. 5-7 were estimated with metabolomics data (**see Table S8 in SI**). As the growth medium contained abundant nutrient sources, there was no constraint on upper bounds of uptake fluxes of phosphate, bicarbonate, choline, and hypoxanthine.

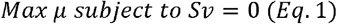

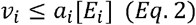

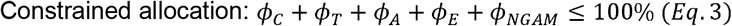

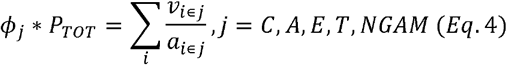

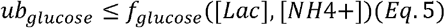

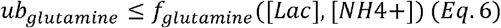

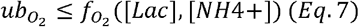

To simulate the growth of *S. scrofa* muscle stem cells dynamically in a batch reactor, dynamic FBA (dFBA) was adopted, as a combination of FBA and differential-equation-based dynamic system modeling (Henson and Hanly, 2014). The concentration changes of extracellular metabolites ([*M*_*i*_]) and *S. scrofa* muscle stem cell biomass ([*X*]) were modeled by differential equations to account for biomass accumulation (Eq. 7) and consumption/production of extracellular metabolites (Eq. 8). Because the growth rate of mammalian cells was relatively low, in comparison to microbial cells, a cell-specific death rate (*k*_*d*_) was needed in dynamic FBA. *k*_*d*_ was set as 0.01/hr in this study (Xu et al., 2010). 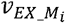 was the exchange flux of the extracellular metabolite *M*_*i*_.

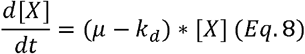

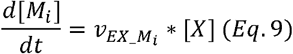

## 3. Results

### 3.1 The effects of lactate and ammonium on *S. scrofa* muscle stem cells

To investigate the effects of accumulating lactate and ammonium, *S. scrofa* muscle stem cells were cultured with two different initial ammonium (NH4+) concentrations (1.073 +/- 0.021 mM and 3.823 +/- 0.015 mM) (**see Table S2&S3 in SI**). The biomass level could accumulate to 0.59 g/L in the low initial NH4+ level condition, which reduced to 0.39 g/L in the high initial NH4+ level condition (**Figure 1A**). For energy metabolism, the glucose and glutamine consumed and lactate produced in high initial NH4+ level condition were approximately half of those in low initial NH4+ level condition (**Figure 1BC**). The NH4+ concentration in the low initial NH4+ level condition was generally below 4mM, while that of high initial NH4+ level condition was generally above 4mM (**Figure 1C**). Therefore, the main inhibitor of cellular growth in the low initial NH4+ level condition was lactate, while the main inhibitor in the high initial NH4+ level condition was NH4+. The computed growth rates of all time intervals showed that the stem cells were not fully activated before 42 hr and 30 hr for the low and high initial NH4+ level conditions (**Figure 1D**). Thereafter, the highest growth rates reached by stem cells in the low and high initial NH4+ conditions were 0.0664/hr and 0.0516/hr, correspondingly. Regarding the inhibitory effects of lactate and NH4+ on the growth rates of *S. scrofa* muscle stem cells, the negative correlations between lactate concentrations and growth rates in both culture conditions were not statistically significant, whereas NH4+ concentrations had significantly negative correlations with growth rates (**Figure 1EF**).

**Figure 1.**
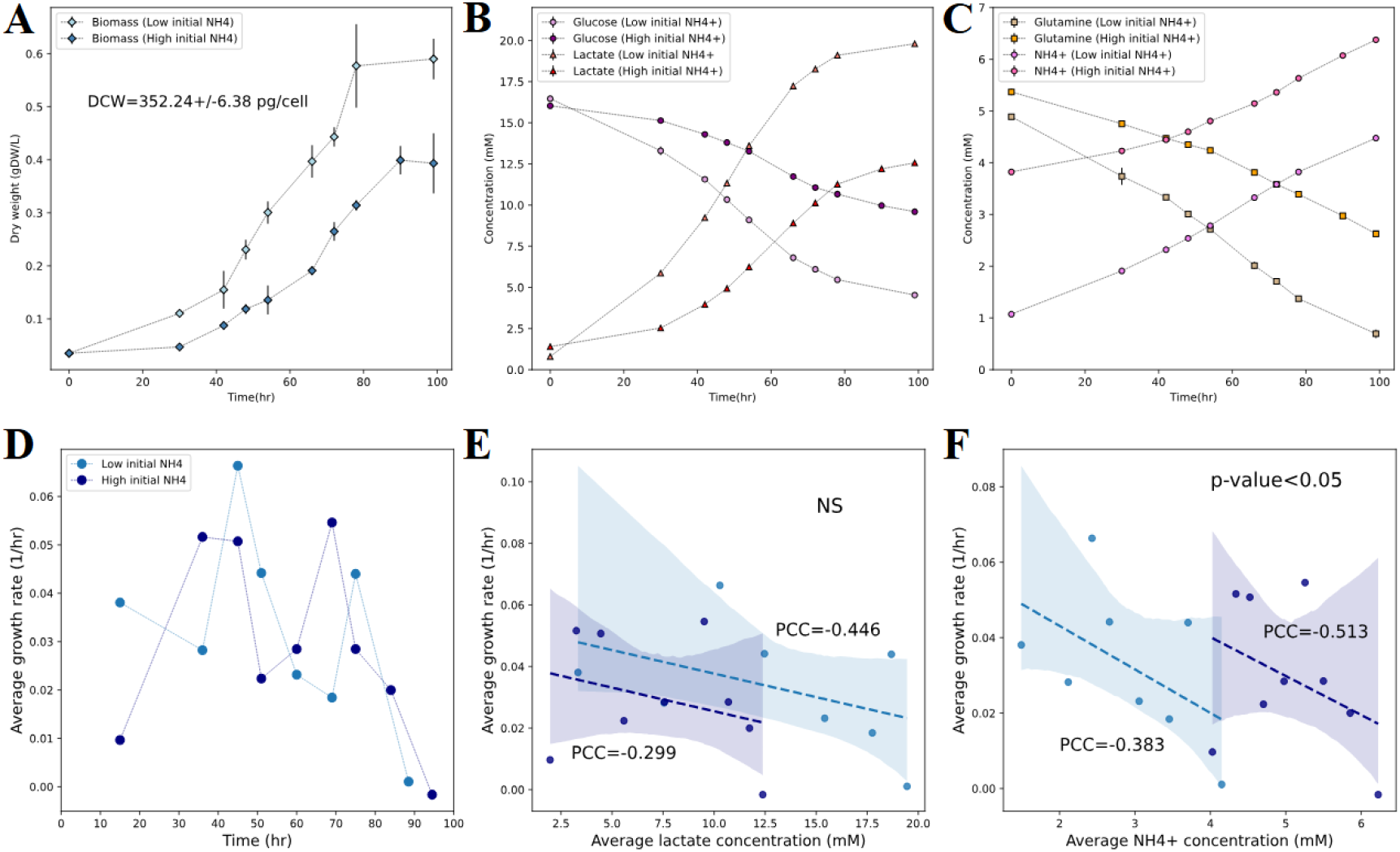
Growth and metabolism of *S. scrofa* muscle stem cells under low and high initial ammonium concentrations (1.073 +/- 0.021 mM and 3.823 +/- 0.015 mM). (A) Dry cellular biomass concentration profiles. DCW: dry cellular weight; (B) Concentration profiles of glucose and lactate. (C) Concentration profiles of glutamine and NH4+. (D) Average growth rates of all time intervals. (E) Pearson correlations between lactate concentrations and growth rates. (F) Pearson correlations between NH4+ concentrations and growth rates.

Cell-specific metabolic fluxes were computed based on biomass and metabolite concentrations to further quantify the metabolic status of *S. scrofa* muscle stem cells (**see section S1**.**1 and Figure S2 in SI**). The highest glucose uptake rates reached in low and high initial NH4+ conditions were 1.45 and 1.0322 mmol/gDW*hr, respectively (**Figure 2A**). In both culture conditions, the lactate:glucose molar ratios were around 1.75 at mid-exponential stage (excluding outliers), and then decreased to below 1.5 at late-exponential stage, indicating a carbon flux redirection from anaerobic fermentation of lactate to aerobic respiration (**Figure 2B**).

**Figure 2.**
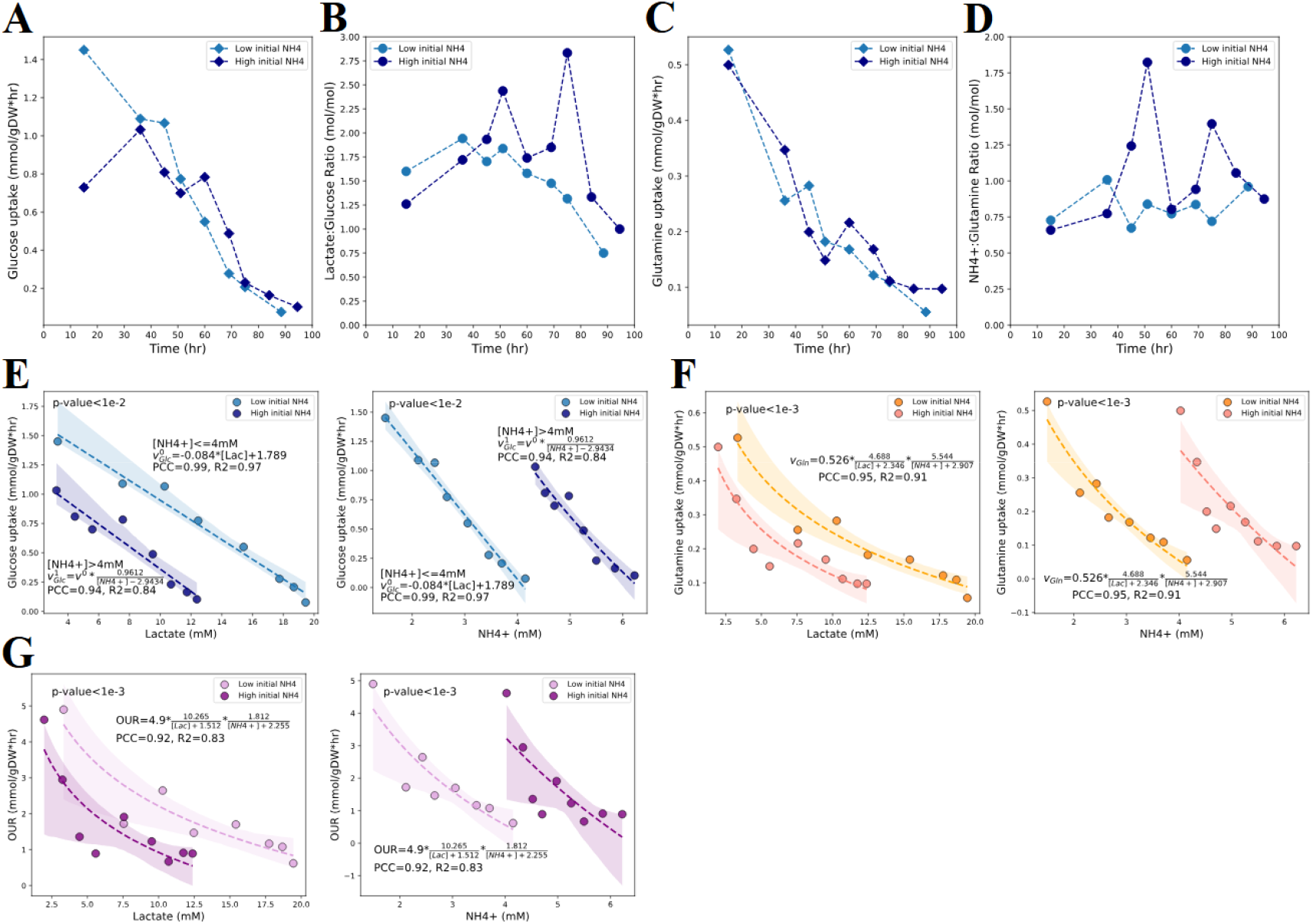
Experimental metabolic fluxes of *S. scrofa* muscle stem cells under low and high initial NH4+ level conditions. (A) Cell-specific glucose uptake rates. (B) Lactate: glucose molar ratios. (C) Cell-specific glutamine uptake rates. (D) NH4+:glutamine molar ratios. (E) The relationship between glucose uptake rates and lactate, NH4+ concentrations (p-value<0.01). (F) The relationship between glutamine uptake rates and lactate, NH4+ concentrations (p-value<0.001). (G) The relationship between oxygen uptake rates (OURs) and lactate, NH4+ concentrations (p-value<0.001).

The redirection of carbon flux from a low proteome cost pathway to a high proteome cost pathway (**Figure 3C**) implied that high lactate and NH4+ concentrations most likely inhibited the expression level of glycolysis rather than enzyme activities, as the reduction of glycolytic enzyme activities would raise the proteome cost of energy metabolism. The lactate:glucose molar ratios of the high initial NH4+ condition were slightly higher than those of the low initial NH4+ condition at mid- and late-exponential phases, suggesting a reduced energy yield from glucose due to the inhibition of oxidative phosphorylation by NH4+ (Ozturk et al., 1992). The highest glutamine uptake rates reached in low and high initial NH4+ conditions were 0.5266 and 0.4995 mmol/gDW*hr, respectively (**Figure 2C**). The NH4+:glutamine molar ratios were mostly fluctuating around 1 in both culture conditions, suggesting that glutamine metabolism was the major source of NH4+ production (**Figure 2D**). Subsequently, the quantitative relationships of lactate and NH4+ concentrations with glucose, glutamine, and oxygen uptake rates (OURs) were approximated based on metabolomics data with empirical functions (**see Table S8 in SI**) to model the inhibitory effects of lactate and NH4+ on glycolysis, amino acid metabolism, and oxidative phosphorylation (Chen et al., 2009; Cruz et al., 2000; Galvanauskas et al., 2019; Ozturk et al., 1992; Roon et al., 1975; Slivac et al., 2010) (**Figure 2E-G**). In short, experimental growth and metabolomics data quantitatively illustrated how accumulated lactate and NH4+ inhibited the growth and metabolism of *S. scrofa* muscle stem cells, contributing to metabolic flux simulation of *S. scrofa* muscle stem cells under varying culture conditions (**section 3**.**3**).

**Figure 3.**
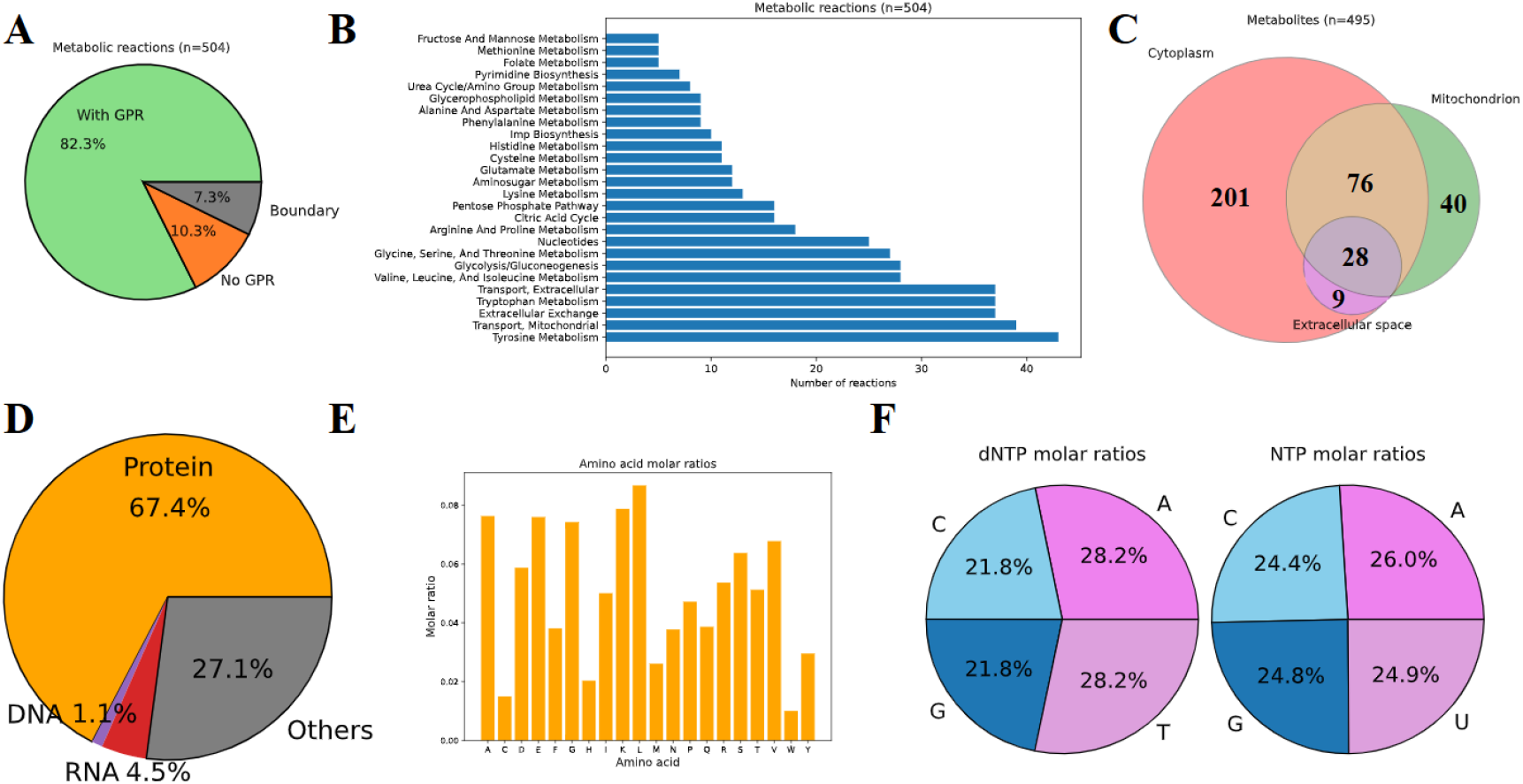
Overview of PigGEM2025. (A) Status of metabolic reaction curation, Green: reactions with GPR rules; Orange: reactions without GPR rules; Grey: exchange reactions at boundary. (B) Summary of curated metabolic reactions in different subsystems. (C) Summary of metabolites in different compartments. (D) Dry biomass composition of the *S. scrofa* stem cell. (E) Molar ratios of amino acids in the proteome of *S. scrofa*. (F) Molar ratios of deoxyribonucleic acids (dNTP) and ribonucleic acids (NTPs) in the genome and transcriptome of *S. scrofa*.

### 3.2 Overview of the genome-scale metabolic model

The reconstructed core GEM of *S. scrofa* muscle stem cells, PigGEM2025, contains 504 reactions and 495 metabolites in total. Among 504 reactions, 82.3% have gene-protein-reaction rules, 7.3% are boundary reactions, and the rest 10.3% miss associated coding genes (**Figure 3A**). The top 5 subsystems (excluding exchange and transport) with most reactions are tyrosine metabolism, tryptophan metabolism, valine/leucine/isoleucine metabolism, glycolysis/gluconeogenesis, and glycine/serine/threonine metabolism (**Figure 3B**). Among 495 metabolites, 314 are in cytoplasm, 144 are in mitochondrion, 37 are in extracellular space (**Figure 3C**). The quality of PigGEM2025 was assessed by MEMOTE (Lieven et al., 2020), and the overall consistency score was 97% (**see Table S4 in SI**). Subsequently, dry biomass composition of the *S. scrofa* muscle stem cell was measured to accurately model the biomass production rate (cellular growth rate) (**see section S1**.**2 in SI**). Proteins, RNA, and DNA take up 67.4%, 4.5%, and 1.1% mass fraction, respectively (**Figure 3D**). Molar ratios of amino acids in *S. scrofa* proteome (Müller et al., 2020) and deoxyribonucleic acids (dNTP) and ribonucleic acids (NTPs) in *S. scrofa* genome and transcriptome (Groenen et al., 2012) were used to estimate mass fractions of 20 essential amino acids, 4 dNTPs, and 4NTPs in dry cellular biomass (**Figure 3EF**). The details of biomass formation reaction can be found in **section S1**.**2 in SI**. To sum up, PigGEM2025 is a GEM tailored for the core metabolism of *S. scrofa* muscle stem cells.

In order to build the proteome constrained FBA model of *S. scrofa* muscle stem cells, pcPigGEM2025 (**section 2**.**4**), enzyme activities were curated for all irreversible reactions (n=272) in PigGEM2025, and 93 out of 272 enzyme activities were experimental data from *S. scrofa* (**Figure 4A**). From curated enzyme activities in different pathways, glycolysis, pentose phosphate pathway, and TCA cycle were more efficient than most amino acid metabolic pathways (e.g., lysine metabolism) and biosynthetic pathways of DNA/RNA building blocks (e.g., pyrimidine biosynthesis) (**Figure 4B**). The integration of proteome costs defined by enzyme activities (**section 2**.**4**) enabled the modeling of lactate overflow at the exponential growth stage of mammalian cell cultures (Young, 2013). The lactate production pathway was much shorter than TCA cycle and oxidative phosphorylation, and the enzyme activity of lactate dehydrogenase (LDH) was higher than most enzymes in TCA cycle (**Figure 4C**). Therefore, due to the limitation of the proteome resource, the mammalian cell, even under aerobic condition, would favor lactate production in place of aerobic respiration to achieve higher metabolic efficiency, despite the greater ATP yield of aerobic respiration. Additionally, the relatively high activity of glutaminase (GLUNm) could explain why the mammalian cell would utilize glutamine as another major carbon source together with glucose (**Figure 4C**). Generally speaking, constrained proteome allocation could improve the performance of FBA by modeling the proteome resource coordination across different metabolic pathways.

**Figure 4.**
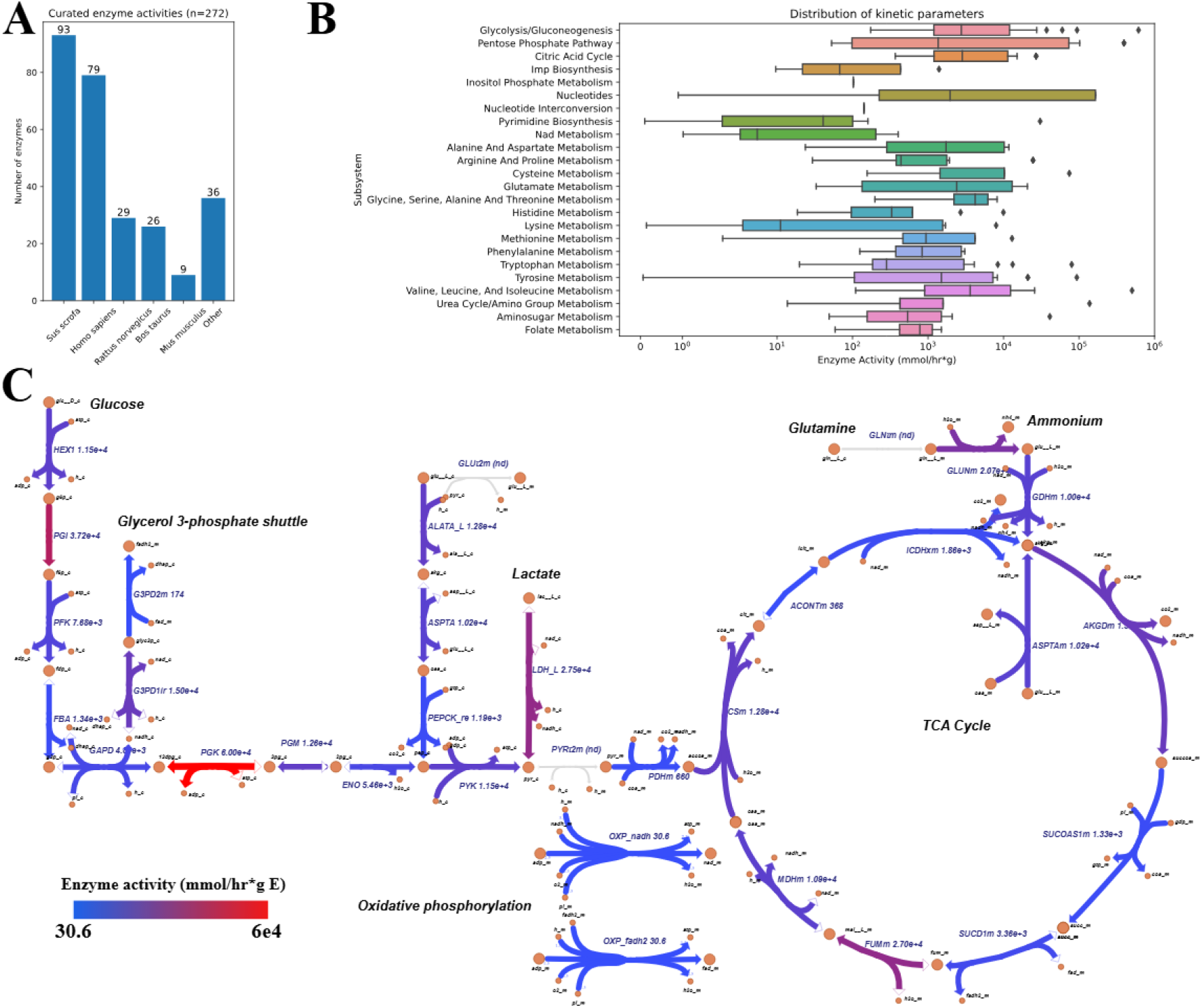
Enzyme specific activities of metabolic reactions in pcPigGEM2025. (A) Enzyme activities curated from *S. scrofa* and mammalian species close to *S. scrofa*. (B) Distribution of enzyme activities of metabolic reactions in different pathways. (C) Enzyme activities (mmol/hr per gram enzyme) mapped to the central carbon metabolic network (full names of reactions can be found in **SI, Table S5**).

### 3.3 Metabolic flux simulation of *S. scrofa* muscle stem cells under varying culture conditions

To examine the prediction performance of pcPigGEM2025, the static metabolic flux simulation was performed for experimental data from selected time intervals in low and high initial NH4+ level conditions (**Figure 5**). Details of experimental data for selected time intervals can be found in **SI, Table S7**. For the low initial NH4+ level condition, the simulation accurately predicted growth rates, glucose consumption rates, and lactate production rates (PCC>0.95, R2>0.7) (**Figure 5ABD**). While the prediction result captured the inhibition of glutamine metabolism by lactate and NH4+ (PCC>0.8), the quantitative accuracy of predicted glutamine consumption and NH4+ production rates was poor (R2<0) (**Figure 5CE**). Compared to the low initial NH4+ level condition, the predicted growth rates at high NH4+ concentrations were less quantitatively accurate (PCC<0.9, R2<0), suggesting that the inhibitory effect of NH4+ was not sufficiently well approximated (**Figure 5AF**). Similar to the low initial NH4+ level condition, the prediction of glucose consumption and lactate production rates was accurate (**Figure 5GI)**, but the prediction of glutamine consumption and NH4+ production rates could only capture the tendency (**Figure 5HJ**). For both low and high initial NH4+ condition, glutamine consumption rates were underestimated and NH4+:glutamine ratios were overestimated (**Figure 5CH**), indicating that the simulation overestimated the NH4+ production from the metabolic pathways of other amino acids (e.g., proline). Like glutamine, proline can also be converted to alpha-ketoglutarate by proline oxidase, 1-pyrroline-5-carboxylate dehydrogenase, and glutamate dehydrogenase (**see Figure S5 in SI**). Expectedly, inaccuracies in static metabolic flux simulations would propagate to dynamic metabolic flux simulations.

**Figure 5.**
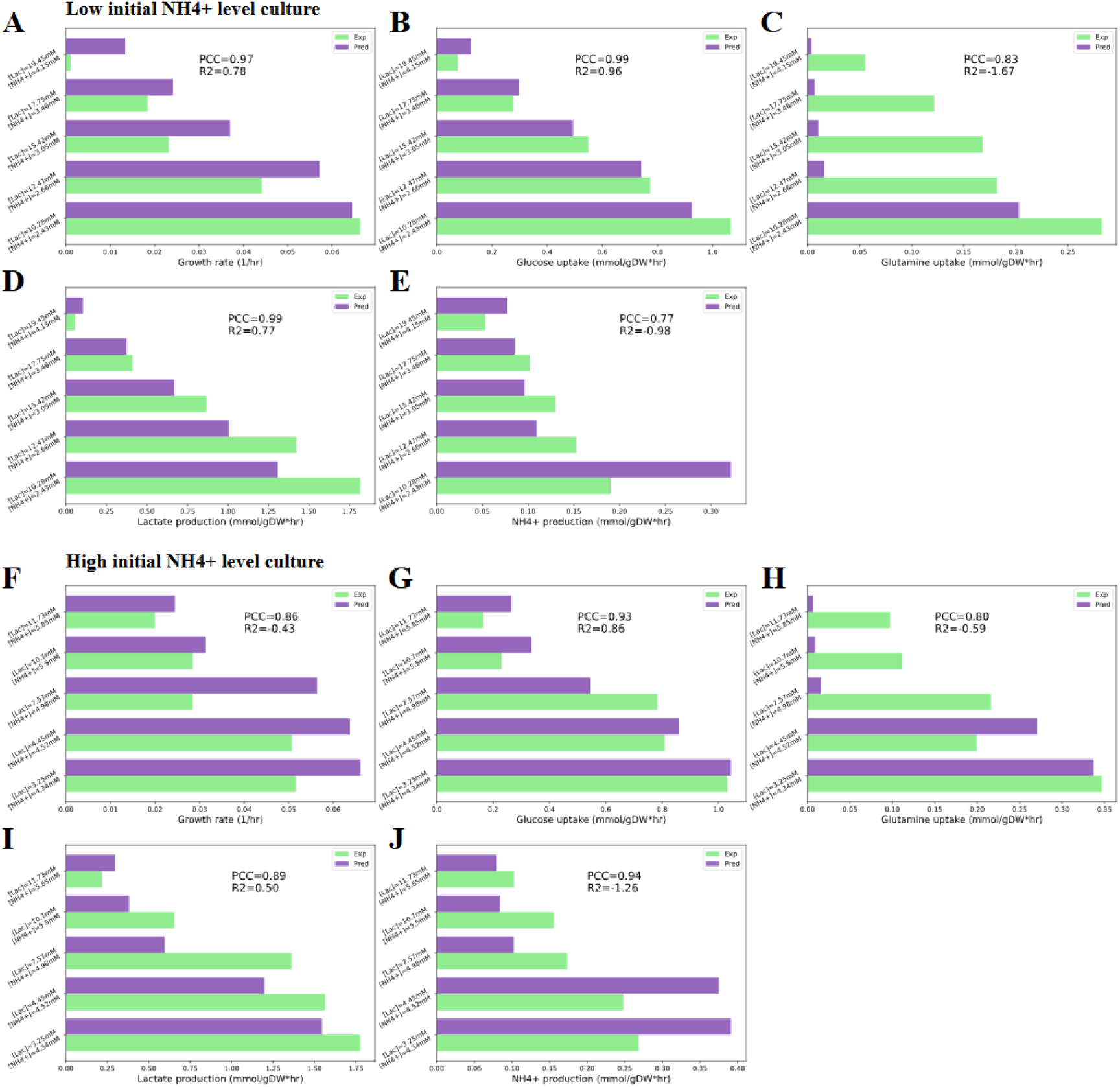
Static metabolic flux simulation for different lactate and NH4+ concentrations. A-E: low initial NH4+ level condition; F-J: high initial NH4+ level condition. (A) Experimental and predicted growth rates under low NH4+ level (Pearson correlation coefficient (PCC)=0.97). (B) Experimental and predicted glucose uptake rates under low NH4+ level (PCC=0.99). (C) Experimental and predicted glutamine uptake rates under low NH4+ level (PCC=0.83). (D) Experimental and predicted lactate production rates under low NH4+ level (PCC=0.99). (E) Experimental and predicted NH4+ production rates under low NH4+ level (PCC=0.77). (F) Experimental and predicted growth rates under high NH4+ level (PCC=0.86). (G) Experimental and predicted glucose uptake rates under high NH4+ level (PCC=0.93). (H) Experimental and predicted glutamine uptake rates under high NH4+ level (PCC=0.80). (I) Experimental and predicted lactate production rates under high NH4+ level (PCC=0.89). (J) Experimental and predicted NH4+ production rates under high NH4+ level (PCC=0.94). Green: experimental data; Purple: prediction results.

The dynamic simulation of *S. scrofa* muscle stem cell proliferation was performed for both low and high initial NH4+ level conditions (**Figure 6**). Two initial conditions were: (1) T=42hr, [glucose]=11.567mM, [glutamine]=3.333mM, [lactate]=9.233mM, [NH4+]=2.320mM; (2) T=30hr, [glucose]=15.133mM, [glutamine]=4.753mM, [lactate]=2.533mM, [NH4+]=4.23mM (**see Table S1&S2 in SI**). For the low initial NH4+ level culture, the prediction result achieved good accuracy for the concentrations of biomass, glucose, lactate, and NH4+, but largely underestimated the consumption rate of glutamine (**Figure 6A-E**). For the high initial NH4+ level culture, the prediction result achieved good accuracy for the concentrations of biomass, glucose, glutamine, and NH4+, but underestimated the production rate of lactate (**Figure 6F-J**). The predicted growth curve of high initial NH4+ level culture was less accurate than that of low initial NH4+ level culture, suggesting that the characterization of the inhibition by NH4+ was not accurate enough (**Figure 6F**). The prediction result of high initial NH4+ level culture also underestimated the consumption rate of glutamine, although the quantitative accuracy was high before T=70hr (**Figure 6H**). In conclusion, pcPigGEM2025 can accurately predict growth kinetics and central carbon metabolism of *S. scrofa* muscle stem cells under varying lactate and NH4+ concentrations, but its quantitative accuracy on amino acid metabolism still requires improvement.

**Figure 6.**
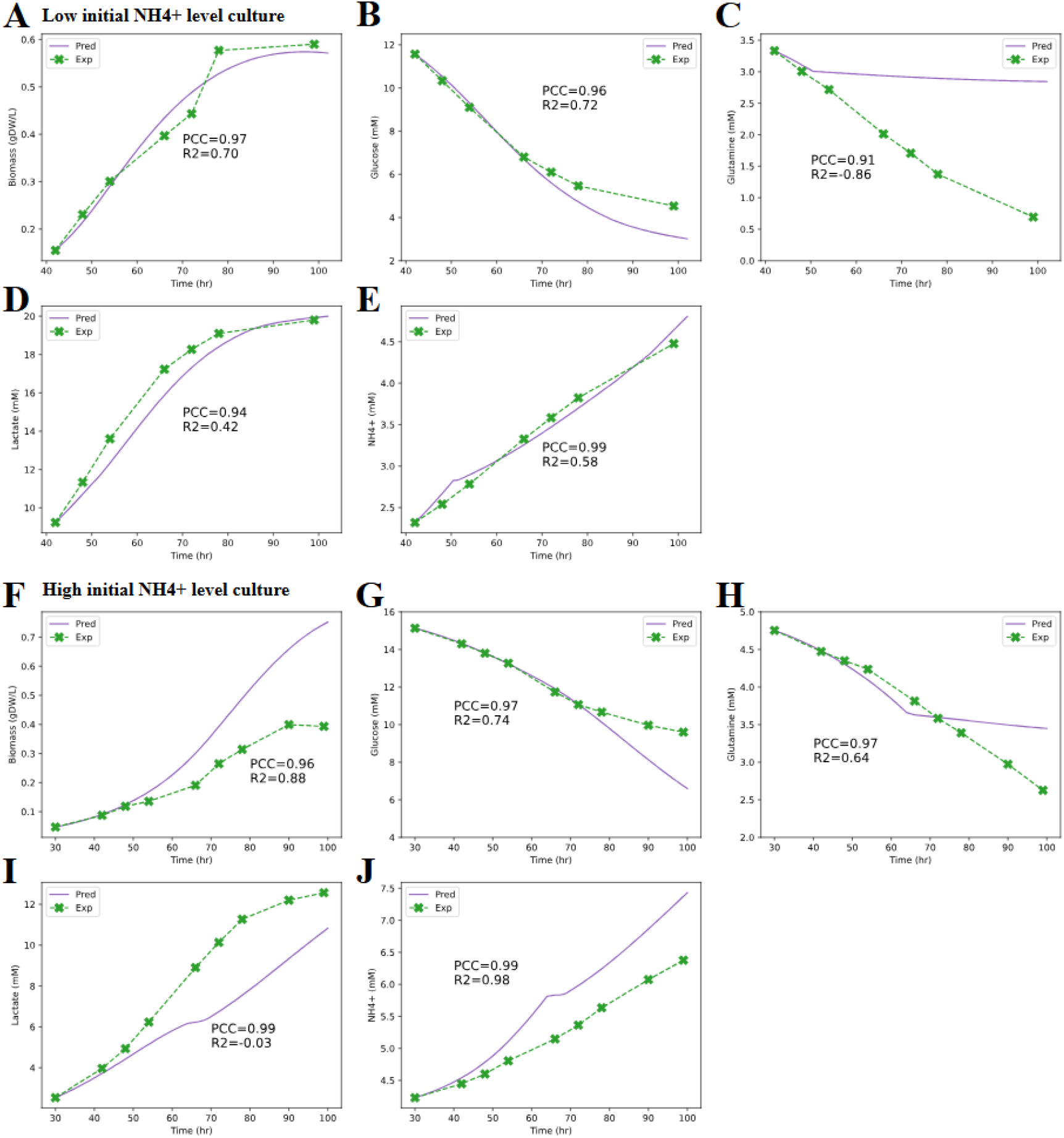
Dynamic simulation of *S. scrofa* muscle stem cell culture with low and high initial NH4+ concentrations. A-E: low initial NH4+ level, from 42 hr to 100 hr; F-J: high initial NH4+ level, from 30 hr to 100 hr. (A) Experimental and predicted biomass concentrations under low NH4+ level (Pearson correlation coefficient (PCC)=0.97). (B) Experimental and predicted glucose concentrations under low NH4+ level (PCC=0.96). (C) Experimental and predicted glutamine concentrations under low NH4+ level (PCC=0.91). (D) Experimental and predicted lactate concentrations under low NH4+ level (PCC=0.94). (E) Experimental and predicted NH4+ concentrations under low NH4+ level (PCC=0.99). (F) Experimental and predicted biomass concentrations under high NH4+ level (PCC=0.96). (G) Experimental and predicted glucose concentrations under high NH4+ level (PCC=0.97). (H) Experimental and predicted glutamine concentrations under high NH4+ level (PCC=0.97). (I) Experimental and predicted lactate concentrations under high NH4+ level (PCC=0.99). (J) Experimental and predicted NH4+ concentrations under high NH4+ level (PCC=0.99). Green: experimental data; Purple: prediction results.

### 3.4 Perturbation simulation for non-growth associated maintenance and oxygen supply

Although the influence of accumulated lactate and NH4+ on NGAM was not observed in this study (**see Figure S3 in SI**), various types of environmental stress (e.g., osmotic pressure) in the bioreactor might affect the NGAM. Usually, the environmental stress will increase the NGAM. Therefore, a perturbation on NGAM values was performed (**Figure 7A-C**). In the simulation, the increase of NGAM led to the decrease of growth rate from 0.0664/hr to 0.0040/hr due to the competition of energy and proteome resources between growth and stress responses (**Figure 7A**). The increase of lactate:glucose molar ratio reflected the inhibition of aerobic respiration, as the carbon flux through anaerobic fermentation of lactate increased (**Figure 7B**). The proteome resource reallocation to the NGAM sector limited the proteome resource available for central carbon metabolism, and thus inhibited aerobic respiration with a higher proteome cost than lactate production (**section 3**.**2**). At high NGAM, the glutamine metabolism was predicted to be the major source of NH4+ production, as glutamine can be converted to a TCA cycle intermediate, alpha-ketoglutarate, with fewer steps than glucose (**Figure 7C**).

**Figure 7.**
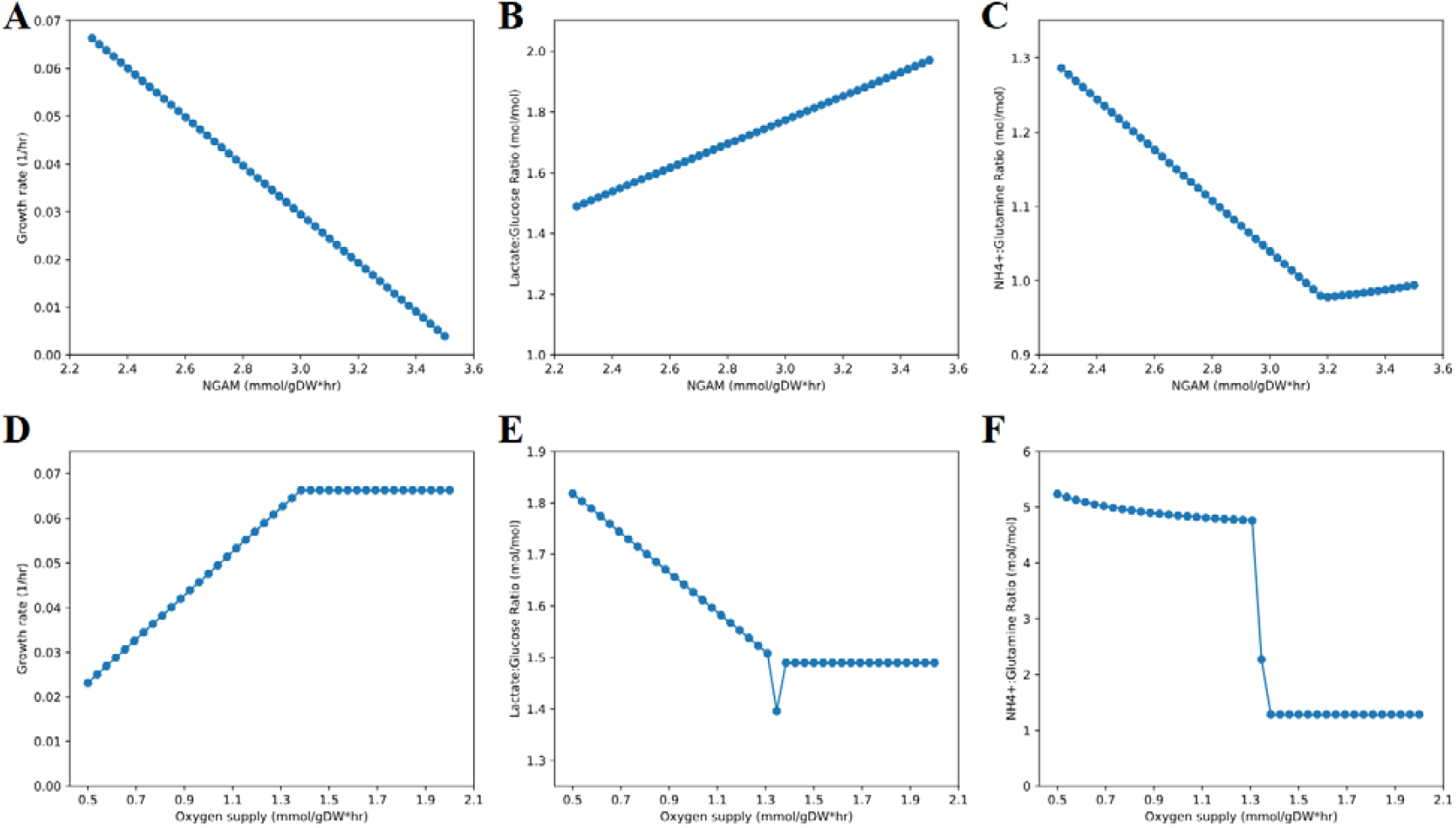
Perturbations on NGAM values and cell-specific oxygen supply levels. (A) Predicted growth rates with different NGAM values. (B) Predicted lactate:glucose molar ratios with different NGAM values. (C) Predicted NH4+:glutamine molar ratios with different NGAM values. (D) Predicted growth rates with different oxygen supply levels. (E) Predicted lactate:glucose molar ratios with different oxygen supply levels. (F) Predicted NH4+:glutamine molar ratios with different oxygen supply levels.

The oxygen supply is a limiting factor of cell proliferation in the bioreactor, and thus, this study also conducted a perturbation on cell-specific oxygen supply (**Figure 7D-F**). The simulation predicted that the increase of oxygen supply would increase the growth rate to 0.0664/hr, the maximum growth rate, and then the growth rate stayed unchanged (**Figure 7D**). The decrease of lactate:glucose ratio indicated the increase of the carbon flux fraction of aerobic respiration (**Figure 7E**), but the lactate:glucose ratio stopped decreasing after oxygen supply reached around 1.4 mmol/gDW*hr due to the limitation of the proteome resource (**section 3**.**2**). The fast drop of NH4+:glutamine ratio suggested that glutamine metabolism became the major source of NH4+ production at high oxygen supply (**Figure 7F**), because glutamine mainly participates in aerobic energy metabolism. In summary, these two perturbation case studies showed that pcPigGEM2025 could also model the effects of other environmental factors, beyond lactate and NH4+ stress.

## 4. Discussion

Given the lack of a computational model for the proliferation of *S. scrofa* stem cells in cultured meat production, this study endeavored to construct a proteome constrained metabolic model named pcPigGEM2025 based on experimental growth and metabolomics data of *S. scrofa* stem cell culture under low and high initial NH4+ levels. The computed growth rates and metabolic fluxes allowed the quantification of the inhibitory effects of accumulated lactate and NH4+ on glycolysis, oxidative phosphorylation, and amino acid metabolism (**section 3**.**1**). Subsequently, pcPigGEM2025, incorporating the effects of lactate and NH4+, adequately modeled the response of *S. scrofa* stem cells’ metabolism to varying lactate and NH4+ concentrations (**section 3**.**3**). The validation with experimental data demonstrated that pcPigGEM2025 could accurately predict growth kinetics, glucose consumption, and lactate secretion. It could effectively capture the response of glutamine consumption and NH4+ secretion to changing concentrations of lactate and NH4+, although the quantitative accuracy remained to be improved. In addition, the perturbation simulation on NGAM and oxygen supply showed that pcPigGEM2025 could also account for other types of environmental stress (**section 3**.**4**). Briefly speaking, this study provided the first metabolic modeling framework for mammalian cells under environmental stress in actual bioprocesses.

Nonetheless, some limitations remained in the presented metabolic model. First, the evaluation of metabolic flux simulation found that the predicted growth rates at high NH4+ concentrations were less accurate than those at low NH4+ concentrations, and the primary error source was that the approximation of NH4+’s inhibitory effect was not accurate enough. Gene expression levels of *S. scrofa* muscle stem cells at different lactate and NH4+ concentrations allow the derivation of mathematical functions that describe proteome resource allocation induced by lactate and NH4+, which may improve the approximation of lactate’s and NH4+’s inhibitory effects (Qiu et al., 2025). Another limitation was the overestimation of NH4+ production from the metabolic pathways of amino acids (e.g., arginine, leucine) other than glutamine, which was most likely caused by the inaccuracy of enzyme activity values (most of them were not specific to *S. scrofa*). The lack of organism-specific kinetic parameters was a well-acknowledged difficulty for proteome constrained metabolic modeling (Nilsson et al., 2017). Deep learning-based models have been developed to predict enzyme kinetic parameters, but the prediction error is not low enough (log10-scale RMSE=∼0.8) to offer accurate parameter estimation (F. Li et al., 2022; Qiu et al., 2023b; Shen et al., 2024). Moreover, the stoichiometric coefficients of biomass components (e.g., 20 essential amino acids) in the objective function (biomass formation) were estimated based on a coarse-grained biomass composition instead of precisely measured, resulting in uncertainty in modeling the growth and metabolism of *S. scrofa* muscle stem cells.

In spite of several limitations discussed above, pcPigGEM2025, presented in this study, exhibited the potential to be a model-aided process engineering tool for cultured meat production, which can be used to design optimal carbon/nitrogen source feeding strategies (Boojari et al., 2023) or co-culture of stem cells and probiotic *E. coli* strains to remove NH4+ (Kolodkin-Gal et al., 2024; Wassenaar, 2016). As envisaged, the accurate simulation of mammalian stem cells’ growth and metabolism in bioreactors will give rise to the digital twin of cultured meat production, benefiting the development of future food.

## Abbreviations

CHO: Chinese hamster ovary
DCW: dry cellular weight
DW: dry weight
GAM: growth associated maintenance
GEM: genome-scale metabolic model
GPR: gene-protein-reaction rule
FBA: flux balance analysis
NGAM: non-growth associated maintenance
NH4+: ammonium
OUR: oxygen uptake rate
TCA: tricarboxylic acid

## Acknowledgements

This work was financially supported by Biotechnology and Biological Sciences Research Council of UK Research and Innovation (project ref. BB/Y007859/1). The authors thank Ivy Farm Technologies (Oxford, UK) and Julia Lee of Ivy Farm Technologies for their technical assistance with the experiments.

## Author contributions

Sizhe Qiu conceptualized the study, developed the methodology, conducted experiments, performed data analysis, and contributed to the writing of the first draft. Eliska Kratochvilova contributed to methodology development, experimental work, and writing. Prof. Wei E. Huang, Prof. Zhanfeng Cui, and Dr. Tom Agnew assisted in the writing and review of the first draft. Prof. Aidong Yang and Prof. Hua Ye supervised this research project and critically reviewed the manuscript.

## Data availability statement

The code and data are openly available at https://github.com/SizheQiu/PigGEM2025 and supplementary materials.

