## Supplementary material for "Proteome constrained metabolic modeling of *Sus scrofa* muscle stem cells for cultured meat production": Figure S1-5, Table S1-8

### Supplementary Information

#### 1. Supplementary methods

##### 1.1 Analysis of growth and metabolomics data

For all time intervals in two different culture conditions (**Table S1&S2**), the average growth rate was calculated using Eq. S1. $[X]$ represents the biomass concentration. $t_1$ and $t_2$ represent two time points. For glucose, glutamine, lactate, and ammonium (NH4+), the cell-specific consumption or production rate was calculated using Eq. S2. $[M]$ represents the concentration of a metabolite. $\frac{[X]_1}{2} + \frac{[X]_2}{2}$ is the average biomass concentration. The oxygen uptake rate (OUR) was not directly measured, but estimated from glucose/glutamine uptake rates and lactate secretion rate (Eq. S3). $v_{glc}$ is glucose uptake rate, $v_{lac}$ is lactate secretion rate, $v_{glc} - \frac{v_{lac}}{2}$ is the flux of glucose utilized in aerobic respiration. $v_{gln}$ is glutamine uptake rate. The aerobic respiration of 1 mole glucose or glutamine will consume 6 moles of oxygen.

$$\mu(t_1 \sim t_2) = \frac{ln(\frac{[X]_2}{[X]_1})}{t_2 - t_1} \ (Eq.\ S1)$$

$$v_M(t_1 \sim t_2) = \frac{[M]_2 - [M]_1}{t_2 - t_1} * \frac{1}{\frac{[X]_1}{2} + \frac{[X]_2}{2}} \ (Eq.\ S2)$$

$$OUR = (v_{glc} - \frac{v_{lac}}{2} + v_{gln}) * 6 \ (Eq.\ S3)$$

Growth associated maintenance (GAM) and non-growth associated maintenance (NGAM) were estimated from the linear relationship of ATP generation rates in energy metabolism ($v_{ATP}$) and growth rates (μ) (Eq. S4). GAM is the slope and NGAM is the intercept on y axis (**Figure S3**). The ATP generation rate was computed with $v_{glc}$, $v_{lac}$, and $v_{gln}$ (Eq. S5). For 1 mole of lactate produced, 1 mole of ATP is generated. For 1 mole of glucose utilized in aerobic respiration, 30 moles of ATP are generated. For 1 mole of glutamine utilized in aerobic respiration, 12.5 moles of ATP are generated.

$$v_{ATP} = GAM * \mu + NGAM \ (Eq.\ S4)$$

$$v_{ATP} = v_{lac} + (v_{glc} - \frac{v_{lac}}{2}) * 30 + v_{gln} * 12.5 \ (Eq.\ S5)$$

#### 1.2 *S. scrofa* muscle stem cell biomass composition measurement

The dry weights per cell of protein, DNA, RNA, and total biomass were measured to define the biomass composition for the *S. scrofa* skeletal myoblasts.

For the determination of total dry biomass, cultures were collected 70 hr after seeding at passage 59 (pre-stationary phase), pelleted at 600 × g for 5 min at room temperature, resuspended in phosphate-buffered saline (PBS; Sigma-Aldrich, UK) supplemented with 0.1 % (w/v) Poloxamer 188 (Sigma-Aldrich, UK), and subjected to one additional wash under the same conditions. Subsequently, cells were resuspended and counted, and three replicates, each containing $1.44 \times 10^{7}$ cells per 570 uL, were transferred into pre-weighed cryovials. Samples were frozen by placing onto the pre-cooled shelf of a freeze dryer (Epsilon 1-4 LSCPlus, Martin Christ, Germany) and lyophilised for at least 48 hr under high vacuum. Vials were equilibrated to room temperature under vacuum before removal, then allowed to equilibrate at ambient conditions prior to weighing on an analytical balance (CP225D, Sartorius, Germany). The dry biomass of each sample was obtained by subtracting the estimated dry weight contribution of the wash buffer, corrected for the volume displaced by cells, from the total dry weight. Biomass per cell was calculated via dividing the corrected biomass by the corresponding cell count.

For protein quantification, cells were harvested at passage 70, approximately 70 hr after seeding, corresponding to the pre-stationary phase. Pellets were obtained by centrifugation at 600 × g for 5 min at 4 °C, washed once in ice-cold PBS (Sigma-Aldrich, UK), and centrifuged again at 1,000 × g for 5 min at 4 °C. The resulting pellets were resuspended in PBS supplemented with Halt™ Protease Inhibitor Cocktail (final 1×; Thermo Fisher Scientific, UK), and cell numbers were determined. Three aliquots of 200 µL were prepared for lysis, and each aliquot contained about $4.7 \times 10^5$ cells. Cells were lysed by three freeze–thaw cycles (liquid nitrogen bath followed by thawing at 37 °C) and incubated on ice for 15 min. Lysates were clarified by centrifugation at 13,000 × g for 10 min at 4 °C. Thereafter, protein concentrations were determined using the QuantiPro™ BCA Assay Kit (Sigma-Aldrich, UK), with samples diluted in PBS to fall within the range of BSA standards (0–30 µg/mL) and measured in triplicates at 562 nm on a SpectraMax i3x Multi-Mode Microplate Reader (Molecular Devices, USA).

Genomic DNA was obtained from cultures harvested 70 hr after seeding at passage 67. Cells were pelleted by centrifugation as described in protein quantification procedure, resuspended in PBS (Sigma-Aldrich, UK), and counted; $4.87 \times 10^6$ cells were used per replicate (three in total). Extraction was carried out with the DNeasy Blood & Tissue Kit (Qiagen, UK). DNA concentration was determined using the Quant-iT™ dsDNA PicoGreen® Assay Kit (Thermo Fisher Scientific, UK), with fluorescence measured across six dilutions per sample. Readings were taken on the same microplate reader as above (excitation ~480 nm, emission ~520 nm).

For RNA quantification, cells were collected 70 hr after seeding at passage 67. Three replicate samples were processed and each contained $3.0 \times 10^6$ cells. Total RNA was purified using the RNeasy Mini Kit (Qiagen, UK) following the manufacturer's protocol, with homogenisation performed using QIAshredder spin columns (Qiagen, UK). RNA yield and purity were assessed on a NanoPhotometer (Implen, Germany), with purity evaluated from A260/280 and A260/230 absorbance ratios.

With quantified biomass composition (**Table S3**), relative molar ratios of amino acids, NTPs, dNTPs, and the estimated GAM value (**Figure S3**), this study formulated the objective function of PigGEM2025 as:

0.396736182 ala__L_c + 0.077871574 cys__L_c + 0.305350584 asp__L_c + 0.395002393 glu__L_c + 0.198216455 phe__L_c + 0.386099129 gly_c + 0.105789533 his__L_c + 0.260340282 ile__L_c + 0.409510381 lys__L_c + 0.450934253 leu__L_c + 0.135823949 met__L_c + 0.196191249 asn__L_c + 0.245357299 pro__L_c + 0.20105409 gln__L_c + 0.279116777 arg__L_c + 0.331490572 ser__L_c + 0.266505554 thr__L_c + 0.352503419 val__L_c + 0.052237587 trp__L_c + 0.153650209 tyr__L_c + 0.006266274 datp_c + 0.00483971 dctp_c + 0.004843721 dgtp_c + 0.006274775 dttp_c + 93.888242453 atp_c + 0.022003735 ctp_c + 0.02234101 gtp_c + 0.022448936 utp_c + 0.018176901 pail_psc_c + 0.130185913 pchol_psc_c + 0.049863661 pe_psc_c + 0.001473803 pglyc_psc_c + 0.016457464 ps_psc_c + 0.02137014 sphmyln_psc_c + 0.0327 g1p_c + 88.66501852753811 h2o_c → 93.8648 adp_c + 93.8648 h_c + 93.8648 pi_c + 0.11246061318535946 ppi_c

The stoichiometric coefficients of lipids (e.g., phosphatidylinositol (pail_psc)) and glucose 1-phosphate (g1p), which represented glycosylation, were adopted from the objective function of the CHOv1 model (Hefzi et al., 2016).

#### 2. Supplementary tables

Table S1. Growth and metabolomics data of low initial NH4+ level condition

| Time (hr) | Cell count (cells/ml) | Glucose (mM) | Glutamine (mM) | Lactate (mM) | NH4+ (mM) |
|---|---|---|---|---|---|
| 0 | 100000.00+/-0.00 | 16.4667+/-0.1155 | 4.8900+/-0.0557 | 0.8000+/-0.0000 | 1.0733+/-0.0208 |
| 30 | 313333.33+/-10408.33 | 13.3000+/-0.2646 | 3.7400+/-0.1646 | 5.8667+/-0.0577 | 1.9100+/-0.0173 |
| 42 | 439666.67+/-100957.09 | 11.5667+/-0.0577 | 3.3333+/-0.0058 | 9.2333+/-0.0577 | 2.3200+/-0.0173 |
| 48 | 654666.67+/-52538.87 | 10.3333+/-0.1155 | 3.0067+/-0.0379 | 11.3333+/-0.0577 | 2.5400+/-0.0265 |
| 54 | 853333.33+/-59651.77 | 9.1000+/-0.0000 | 2.7167+/-0.0416 | 13.6000+/-0.1732 | 2.7833+/-0.0404 |
| 66 | 1126666.67+/-87368.95 | 6.8000+/-0.1000 | 2.0133+/-0.0702 | 17.2333+/-0.1528 | 3.3267+/-0.0252 |
| 72 | 1258333.33+/-52041.65 | 6.1000+/-0.0000 | 1.7067+/-0.0252 | 18.2667+/-0.0577 | 3.5833+/-0.0058 |
| 78 | 1638333.33+/-223960.56 | 5.4667+/-0.0577 | 1.3733+/-0.0404 | 19.1000+/-0.2000 | 3.8233+/-0.0321 |
| 99 | 1675000.00+/-108972.47 | 4.5333+/-0.1155 | 0.6933+/-0.0833 | 19.8000+/-0.0000 | 4.4767+/-0.0231 |

| Table S2. Growth and metabolomics data of high initial NH4+ level condition | | | | | |
|---|---|---|---|---|---|
| Time (hr) | Cell count (cells/ml) | Glucose (mM) | Glutamine (mM) | Lactate (mM) | NH4+ (mM) |
| 0 | 100000.00+/-0.00 | 16.0333+/-0.1528 | 5.3700+/-0.0656 | 1.4000+/-0.0000 | 3.8233+/-0.0153 |
| 30 | 133666.67+/-9018.50 | 15.1333+/-0.1528 | 4.7533+/-0.0058 | 2.5333+/-0.0577 | 4.2300+/-0.0346 |
| 42 | 248333.33+/-12583.06 | 14.3000+/-0.1000 | 4.4733+/-0.0577 | 3.9667+/-0.0577 | 4.4467+/-0.0289 |
| 48 | 336666.67+/-25658.01 | 13.8000+/-0.1000 | 4.3500+/-0.0300 | 4.9333+/-0.0577 | 4.6000+/-0.0300 |
| 54 | 385000.00+/-77620.87 | 13.2667+/-0.0577 | 4.2367+/-0.0115 | 6.2333+/-0.0577 | 4.8067+/-0.0153 |
| 66 | 541666.67+/-23094.01 | 11.7333+/-0.1528 | 3.8133+/-0.0603 | 8.9000+/-0.0000 | 5.1467+/-0.0379 |
| 72 | 751666.67+/-50083.26 | 11.0667+/-0.1528 | 3.5833+/-0.0321 | 10.1333+/-0.0577 | 5.3633+/-0.0321 |
| 78 | 891666.67+/-28431.20 | 10.6667+/-0.1155 | 3.3900+/-0.0600 | 11.2667+/-0.1155 | 5.6333+/-0.0153 |
| 90 | 1133333.33+/-76376.26 | 9.9667+/-0.1155 | 2.9733+/-0.0569 | 12.2000+/-0.1732 | 6.0733+/-0.0058 |
| 99 | 1116666.67+/-160727.51 | 9.6000+/-0.1000 | 2.6267+/-0.0569 | 12.5667+/-0.0577 | 6.3767+/-0.0321 |

| Table S3. Biomass composition of *S. scrofa* muscle stem cell | |
|---|---|
| Biomass component | Dry mass (pg/cell) |
| Protein | 237.29+/-11.35 |
| DNA | 3.78+/-0.03 |
| RNA | 15.75+/-0.39 |
| Total biomass | 352.24+/-6.38 |

| Table S4. Model consistency assessment of PigGEM2025 | |
|---|---|
| Stoichiometric Consistency | 100.0% |
| Mass Balance | 98.7% |
| Charge Balance | 98.7% |
| Metabolite Connectivity | 98.2% |
| Unbounded Flux In Default Medium | 86.3% |
| Total score | 97% |

| Table S5. Metabolic reaction information | | | |
|---|---|---|---|
| ID | Name | Compartment | EC number |
| HEX | Hexokinase (D-glucose:ATP) | c | 2.7.1.2 |
| PGI | Glucose-6-phosphate isomerase | c | 5.3.1.9 |
| PFK | Phosphofructokinase | c | 2.7.1.11 |
| FBA | Fructose-bisphosphate aldolase | c | 4.1.2.13 |
| TPI | Triose-phosphate isomerase | c | 5.3.1.1 |
| GAPD | Glyceraldehyde-3-phosphate dehydrogenase | c | 1.2.1.12 |
| PGK | Phosphoglycerate kinase | c | 2.7.2.3 |
| PGM | Phosphoglycerate mutase | c | 5.4.2.11 |
| ENO | Enolase | c | 4.2.1.11 |
| PYK | Pyruvate kinase | c | 2.7.1.40 |
| LDH | Lactate dehydrogenase | c | 1.1.1.27 |
| PDHm | Pyruvate dehydrogenase | m | 1.2.7.1 |
| CSm | Citrate synthase | m | 2.3.3.16 |
| ACONTm | Aconitate hydratase | m | 4.2.1.3 |
| ICDHxm | Isocitrate dehydrogenase | m | 1.1.1.41 |
| GLUNm | Glutaminase (mitochondrial) | m | 3.5.1.2 |
| GDHm | Glutamate dehydrogenase (mitochondrial) | m | 1.4.1.2 |
| ASPTAm | Aspartate transaminase (mitochondrial) | m | 2.6.1.1 |
| AKGDm | 2-oxoglutarate dehydrogenase | m | 1.2.4.2 |
| SUCOAS1m | Succinyl-CoA ligase (GDP-forming) | m | 6.2.1.4 |
| SUCD1m | Succinate dehydrogenase | m | 1.3.5.1 |
| FUMm | Fumarase (mitochondrial) | m | 4.2.1.2 |
| MDHm | Malate dehydrogenase (mitochondrial) | m | 1.1.1.37 |
| PEPCK_re | Phosphoenolpyruvate carboxykinase (GTP) | c | 4.1.1.32 |

| | | | |
|---|---|---|---|
| ASPTA | Aspartate transaminase (cytoplasm) | c | 2.6.1.1 |
| ALATA_L | L-alanine transaminase | c | 2.6.1.2 |
| G3PD1ir | Glycerol 3 phosphate dehydrogenase (NAD) | c | 1.1.1.94 |
| G3PD2m | Glycerol-3-phosphate dehydrogenase (FAD, mitochondrial) | m | 1.1.99.5 |
| OXP_nadh | Oxidative phosphorylation (NADH) | m | 7.1.1.9/7.2.2.19 |
| OXP_fadh2 | Oxidative phosphorylation (FADH2) | m | 7.1.1.9/7.2.2.19 |

* Detailed enzyme and reaction information can be found in the BIGG database (King et al., 2016).

** c: cytoplasm, m: mitochondrion

| Table S6. Reaction enzyme specific activity values* | | | |
|---|---|---|---|
| Reaction | $a_i$ ($\frac{mmol}{hr * g\ E}$) | Organism | Reference |
| EX_glc__D_e | 7.2 | Homo sapiens | (Doege et al., 2001) |
| EX_lac__L_e | 6360 | Saccharomyces cerevisiae | (Schumacher, 2018) |
| EX_aa_e (exchanges of amino acids) | 7.2 | _ | Estimated |
| LDH | 27510 | Homo sapiens | (Pettit et al., 1981) |
| PDHm | 660 | Sus scrofa | (Hamada et al., 1976) |
| CSm | 12780 | Saccharomyces cerevisiae | (Pettersson et al., 2000) |
| ACONTm | 367.8 | Rattus rattus | (Eprintsev et al., 2002) |
| ICDHxm | 1860 | Sus scrofa | (Ramachandran and Colman, 1978) |

| | | | |
|---|---|---|---|
| GLUNm | 20700 | Rattus norvegicus | (Haser et al., 1985) |
| GDHm | 10020 | Bos taurus | (Cho et al., 1995) |
| ASPTAm | 10200 | Sus scrofa | (Barra et al., 1976) |
| AKGDm | 15120 | Urocitellus richardsonii | (Green and Storey, 2020) |
| SUCOAS1m | 1326 | Columba livia | (Johnson et al., 1998) |
| SUCD1m | 3360 | Homo sapiens | (Ziegler and Rieske, 1967) |
| FUMm | 27000 | Sus scrofa | (Sacchettini et al., 1986) |
| MDHm | 10920 | Sus scrofa | (Keighron and Keating, 2010) |
| OXP_nadh/OXP_fadh2 | 30.6 | Mus musculus | (Abe et al., 2005) |
| BIOMASS (Growth) | 107.4 | _ | (Regueira et al., 2021) |
| ATPM (NGAM) | 5.6762 | _ | Estimated |

* Other enzyme specific activity values can be found in

https://github.com/SizheQiu/PigGEM2025/blob/main/data/gems/EnzymeActivity.csv.

Table S7. Experimental data used in metabolic flux simulation and validation

| Condition | Interval | Lactate (mM) | NH4+ (mM) | μ (1/hr) | GlcUR ($\frac{mmol}{hr * gDW}$) | GlnUR ($\frac{mmol}{hr * gDW}$) | OUR ($\frac{mmol}{hr * gDW}$) |
|---|---|---|---|---|---|---|---|
| Low initial NH4+ | 42hr~48hr | 10.28 | 2.43 | 0.0664 | 1.0665 | 0.2825 | 2.65 |
| Low initial NH4+ | 48hr~54hr | 12.47 | 2.66 | 0.0442 | 0.7740 | 0.1820 | 1.47 |
| Low initial NH4+ | 54hr~66hr | 15.42 | 3.06 | 0.0232 | 0.5496 | 0.1681 | 1.70 |
| Low initial NH4+ | 66hr~72hr | 17.75 | 3.46 | 0.0184 | 0.2778 | 0.1217 | 1.17 |

| | | | | | | | |
|---|---|---|---|---|---|---|---|
| Low initial NH4+ | 78hr~99hr | 19.45 | 4.15 | 0.0011 | 0.0762 | 0.0555 | 0.62 |
| High initial NH4+ | 30hr~42hr | 3.25 | 4.34 | 0.0516 | 1.0322 | 0.3468 | 2.95 |
| High initial NH4+ | 42hr~48hr | 4.45 | 4.52 | 0.0507 | 0.8088 | 0.1995 | 1.36 |
| High initial NH4+ | 54hr~66hr | 7.57 | 4.98 | 0.0285 | 0.7829 | 0.2162 | 1.91 |
| High initial NH4+ | 72hr~78hr | 10.70 | 5.50 | 0.0285 | 0.2303 | 0.1113 | 0.67 |
| High initial NH4+ | 78hr~90hr | 11.73 | 5.85 | 0.0200 | 0.1636 | 0.0974 | 0.91 |

Table S8. Approximation performances of empirical mathematical forms for Eq. 5-7

| Equation | Mathematical forms | Parameters | R2 |
|---|---|---|---|
| Eq. 5 | $k_1 * [Lac] + k_2\ if\ [NH4+] \leq 4$ <br> $Else,\ (k_1 * [Lac] + k_2) * \frac{k_3}{[NH4+]+k_4}$ | $k_1 = -0.0841$ <br> $k_2 = 1.7890$ <br> $k_3 = 0.9612$ <br> $k_4 = -2.9434$ | 0.97/0.84 |
| Eq. 5 | $v_{max} * e^{-k_1*([Lac]+k_2)} * e^{-k_3*([NH4+]+k_4)}$ | $v_{max} = 1.45$ <br> $k_1 = 0.0766$ <br> $k_2 = 36.1794$ <br> $k_3 = 0.2072$ <br> $k_4 = 10.0600$ | 0.81 |
| Eq. 5 | $v_{max} * \frac{k_1}{[Lac]+k_2} * \frac{k_3}{[NH4+]+k_4}$ | $v_{max} = 1.45$ <br> $k_1 = 6.1436$ <br> $k_2 = 5.9007$ <br> $k_3 = 6.0443$ <br> $k_4 = 2.0889$ | 0.69 |
| Eq. 6 | $v_{max} * \frac{k_1}{[Lac]+k_2} * \frac{k_3}{[NH4+]+k_4}$ | $v_{max} = 0.526$ <br> $k_1 = 4.6877$ <br> $k_2 = 2.3459$ | 0.91 |

| | | | |
|---|---|---|---|
| | | $k_3 = 5.5442$<br>$k_4 = 2.9073$ | |
| Eq. 7 | $v_{max} * \frac{k_1}{[Lac]+k_2} * \frac{k_3}{[NH4+]+k_4}$ | $v_{max} = 4.9$<br>$k_1 = 10.2652$<br>$k_2 = 1.5119$<br>$k_3 = 1.8122$<br>$k_4 = 2.2553$ | 0.83 |

### 3. Supplementary figures

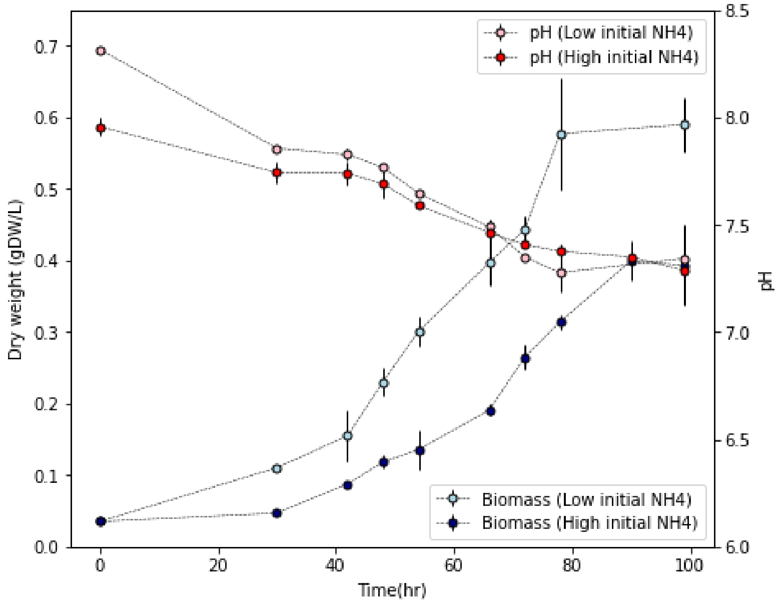


Figure S1. pH recordings for two culture conditions

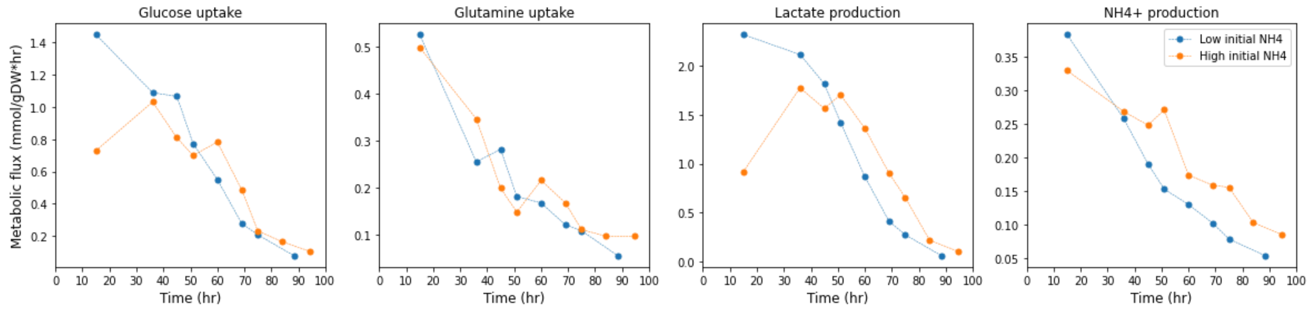


Figure S2. Computed metabolic fluxes of glucose uptake, glutamine uptake, lactate production, and NH4+ production (left to right).

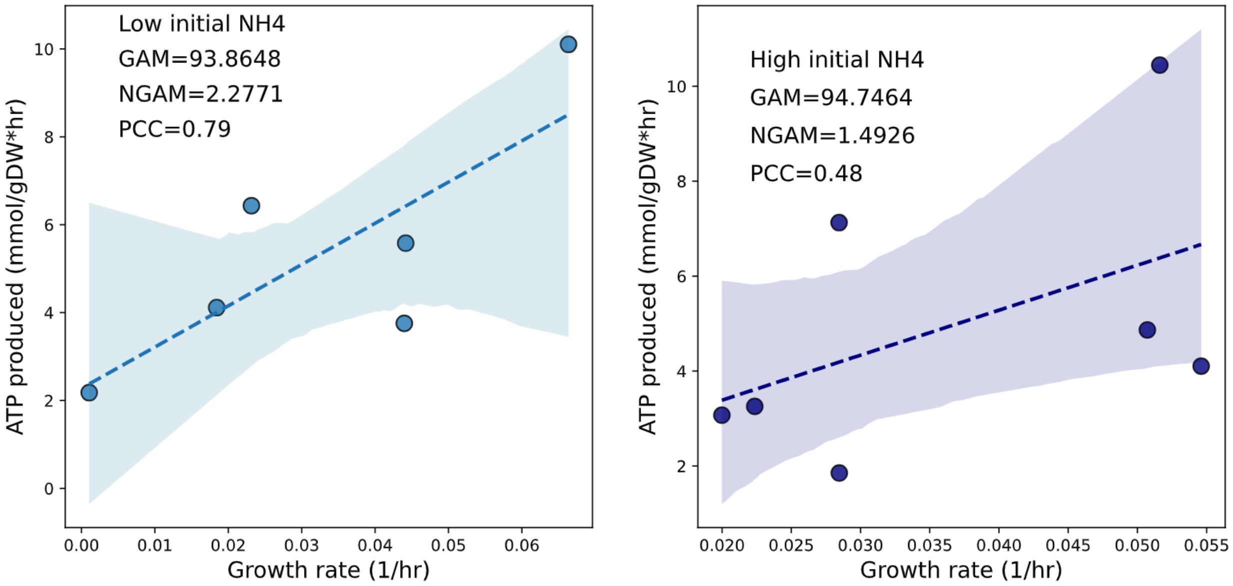


Figure S3. GAM and NGAM values of S. scrofa muscle stem cells under low and high initial NH4+ levels

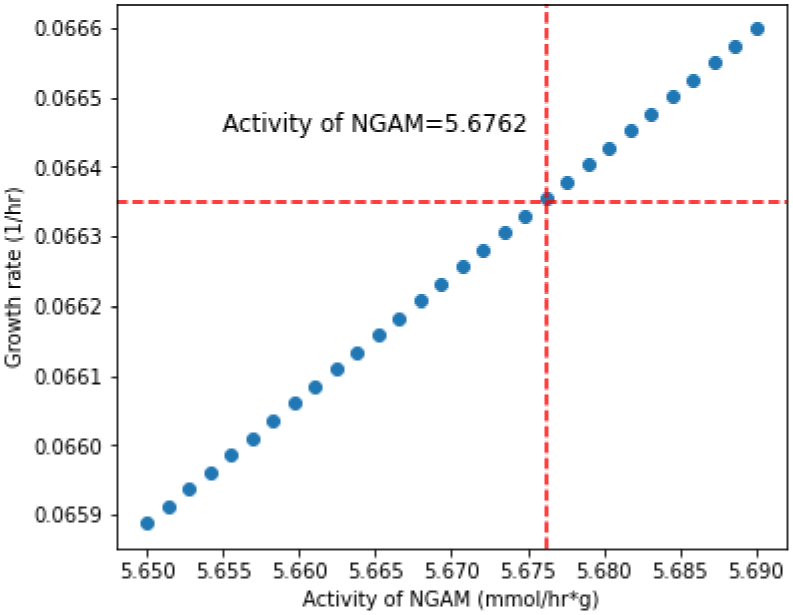


Figure S4. Estimation of the activity value of the NGAM sector. Maximum growth rate=0.6635/hr.

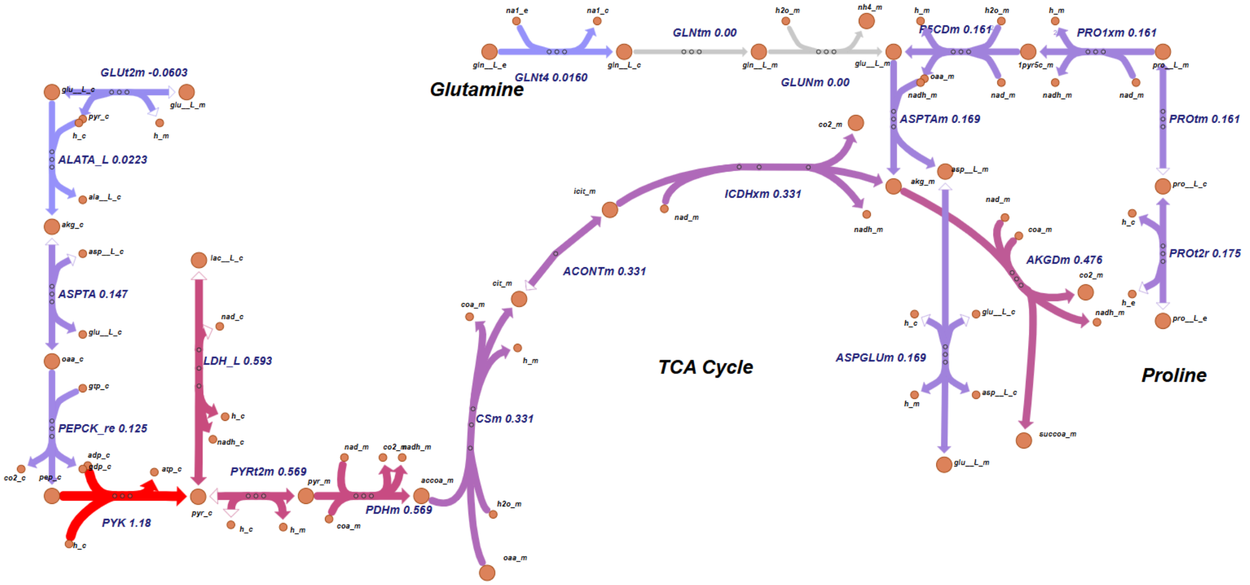


Figure S5. Simulated metabolic fluxes of glutamine and proline metabolic pathways at lactate concentration=7.57mM and NH4+ concentration=4.98mM.
